# Assessment of Flourishing Levels of Individuals by Using Resting State fNIRS with Different Functional Connectivity Measures

**DOI:** 10.1101/2021.01.18.427167

**Authors:** Aykut Eken

## Abstract

Flourishing is an important criterion to assess wellbeing, however, controversies remain, particularly around assessing it with self-report measures. Due to this reason, to be able to understand the underlying neural mechanisms of well-being, researchers often utilize neuroimaging techniques. However, rather than individual answers, previous neuroimaging studies using statistical approaches provided an answer in average sense. To overcome these problems, we applied machine learning techniques to discriminate 43 highly flourishing from regular flourishing individuals by using a publicly available resting state functional near infrared spectroscopy (rs-fNIRS) dataset to get an answer in individual level. We utilized both Pearson’s correlation (CC) and Dynamic Time Warping (DTW) algorithm to estimate functional connectivity from rs-fNIRS data on temporo-parieto-occipital region as input to nine different machine learning algorithms. Our results revealed that by utilizing oxyhemoglobin concentration change with Pearson’s correlation (CC – ΔHbO) and deoxy hemoglobin concentration change with dynamic time warping (DTW – ΔHb), we could be able to classify flourishing individuals with 90 % accuracy with AUC 0.90 and 0.93 using nearest neighbor and Radial Basis Kernel Support Vector Machine. This finding suggests that temporo-parieto-occipital regional based resting state connectivity might be a potential biomarker to identify the levels of flourishing and using both connectivity measures might allow us to find different potential biomarkers.

## 1. Introduction

Well-being is one of the most critical public health issues in populations (Moore et al., 2018). Although there is no consensus about its definition, World Health Organization (WHO) defines well-being for an individual as a self-realization of potential, ability to handle the stress in life, ability to work efficiently and adding values to the community (WHO, 2014). It has been extensively studied and also reviewed for different populations such as mothers (Alderdice et al., 2013), entrepreneurs (Stephan, 2018) and sportsman (Breslin et al., 2017). Well-being is used as important indicator of mental health and the main assumption that lies behind it is; “well-being would prevail when pathology was absent” (Huppert & So, 2013).

Understanding well-being level of individuals is a challenging problem due to being dependent to self-reporting which is a subjective criteria (Diener, 1984). To objectively assess well-being, several neuroimaging studies have been performed to understand the neural correlates of well-being and it was reported that anterior cingulate cortex (ACC), orbitofrontal cortex (OFC), posterior cingulate cortex (PCC), superior temporal gyrus (STG) and thalamus was strongly associated to well-being and these regions are sub-components of default mode network (DMN) (see review (King, 2019)). This finding makes resting state functional connectivity (rsFC) based studies using fMRI (Kong, Liu, et al., 2015; Kong, Wang, et al., 2015; Kong, Wang, et al., 2016; Kong, Xue, et al., 2016; Luo et al., 2014; Luo et al., 2016; Luo et al., 2017; W. Sato et al., 2019) and fNIRS (F. Goldbeck et al., 2018) important to understand the underlying neural mechanisms of well-being.

In this study, we utilized a previously collected rs-fNIRS dataset (F. Goldbeck, 2018) and machine learning techniques to classify the individuals according to the levels of flourishing measured by a flourishing scale. To assess well-being, flourishing scale is a popular measure and widely used in well-being studies (Schotanus-Dijkstra et al., 2016) and flourishing is a term that describes “living within an optimal range of human functioning” (Fredrickson & Losada, 2005) and it is considered as a presence of mental health (Keyes, 2002). On the other hand, fNIRS has recently emerged as an alternative neuroimaging tool to conventional neuroimaging approaches such as fMRI, EEG etc. Main advantages of fNIRS that led to become popular among researchers are; being non-invasive, easy to use, less prone to head motion, low-cost, portability, having high temporal resolution. Since 1990s, it has widely been preferred by researchers from many different fields including neurodevelopment, cognition, anesthesia, psychiatric disorders etc. (see review (Boas et al., 2014)). rs-fNIRS studies has been conducted (see review (Niu & He, 2014)) and several studies validated efficiency of fNIRS to reveal resting state networks by performing simultaneous fMRI and fNIRS measurement (Duan et al., 2012; Sasai et al., 2012; White et al., 2009). Also, rs-fNIRS provides several meaningful inputs to machine learning models to predict mood (Fukuda et al., 2014; M. Sato et al., 2013) or classify psychiatric or neurological disorders (Cheng et al., 2019; J. Li et al., 2016). Also, while estimating the connectivity matrices we utilized the conventional Pearson’s correlation coefficient (CC) and Dynamic Time Warping (DTW) distance. DTW is an elastic matching algorithm that allows to capture the lags between two time series and has recently been used as a functional connectivity metric in several neuroimaging studies (Gokcay et al., 2019; Jin et al., 2020; Linke et al., 2020; Meszlényi et al., 2016; Meszlenyi, Buza, et al., 2017; Meszlenyi, Hermann, et al., 2017; Mohanty et al., 2020).

From machine learning perspective, prediction of well-being was carried out by using Gradient Boosting classifier on 10518 Chinese adolescents via an online survey and subjects’ subjective well-being was predicted with %90 accuracy (N. Zhang et al., 2019). Using domotic sensor data and random forest classifier, mean absolute error of prediction of physical, mental and general health index of elderly individuals were found as %32, %13 and %17, respectively (Casaccia et al., 2020). Other measures such as physiological data (heart rate, skin conductance), phone, mobility and modifiable behavior features were also used in a previous study and classification of high or low stress, high or low mental health, and high or low stress groups was found as 78%, 86% and %74 respectively (Sano et al., 2018). Another study that utilizes multimodal data such as laboratory, demographic and lifestyle to predict wellness found the highest AUC value as 0.726 by using machine learning algorithms such as SVM, Bagging, Adaboost, Random Forest (Agarwal et al., 2016).

In line of these information, our primary objective is to develop a highly accurate model that allows us to classify highly and regular flourishing individuals and also to reveal the potential neural markers that cause this discrimination by utilizing functional connections from individuals. To our best knowledge, this is the first study that uses neuroimaging data to classify flourishing levels and our research question was “Can we find potential biomarkers that allow us to classify individuals according to their levels of flourishing by using resting state fNIRS data and ML techniques?”. For prediction of well-being and conventional statistical approaches used for both self-reporting scales and neuroimaging data provide us an answer on average sense which might be misleading for interpretation of a biomarker for individual prediction of subjective well-being. Classifying highly flourishing individuals is a crucial problem because it will be possible to identify potential objective biomarkers that might lead new perspective in neural correlates of well-being research.

## 2. Methods

In this study, we used a publicly available RS-fNIRS dataset (F. Goldbeck, 2018) collected from 43 individuals who were identified as highly flourishing (HF) and normal flourishing (NF). Details of the subject recruitment, ethical approval, demographic and experimental information can be found in (F. Goldbeck et al., 2018). Participants were sub-grouped as HF / NF by using Flourishing Scale (Diener et al., 2010; Esch et al., 2013). In flourishing scale, participants having scores higher than 48 were assigned to HF and the rest was assigned to NF group. This threshold was determined by using a median-split approach to create a balanced dataset (F. Goldbeck et al., 2018). By using this approach, among 43 participants 23 of them and 20 of them were assigned to NF and HF groups, respectively. In addition to flourishing scale, trait rumination was tested by utilizing the subscale rumination of ruminative response scale (RRS) (Nolen-Hoeksema & Morrow, 1991). Also, depression module of patient health questionnaire (PHQ -9) (Kroenke et al., 2001) was used to assess depressive symptoms and open-thought protocol (OTP) was applied to participants to evaluate their personal experiences during the experiment (Rosenbaum et al., 2017).

### 2.1. fNIRS System

In this study, fNIRS measurements were carried out by using a Hitachi ETG-4000 continuous wave (CW) fNIRS system (Hitachi Co. Japan). The system uses near infrared (NIR) light both in 695 and 830 nm wavelengths and NIR light was sent via a source optode and emitted by using a detector optode. A 3 x 11 optode configuration that includes 17 sources and 16 detectors was used and every source-detector (SD) couple was considered as a channel. In this configuration, 52 channels were used to collect fNIRS time series. SD separation distance were 3 cm and no short SD channel was used to obtain superficial (scalp) signals to regress out for further data analysis. Optical intensity data was converted to oxyhemoglobin (ΔHbO) and deoxyhemoglobin (ΔHb) concentration changes by using Modified Beer-Lambert law (Cope & Delpy, 1988). Sampling frequency of the system was 10 Hz.

### 2.2. Probe Positioning

Optodes were located onto temporo-parieto-occipital region. According to the EEG 10-20 system (Jasper, 1958) T3, T4 and Pz channels were selected as reference points. Previous studies revealed that default mode network (DMN) comprises regions such as precuneus, temporoparietal junction (Raichle et al., 2001) and angular gyrus (Igelstrom & Graziano, 2017). Also, covered regions were reported in previous studies that they have significant associations (see review (King, 2019)). Channel coordinates were obtained by using a navigation system on a subject’s head. Channel positions and corresponding regions were reported in Table 1. Located regions are primary somatosensory cortex (PSC - BA 3,1,2), Supramarginal Gyrus (SupG - BA 40), Superior Temporal Gyrus (STG - BA 22), Middle Temporal Gyrus (MTG - BA 21), Angular Gyrus (AngG - BA 39), Somatosensory Association Cortex (SAC - BA 7) and Third Visual Cortex (V3 – BA 19), Fusiform Gyrus (FusG – BA 37) and Subcentral Area (SC – BA 43). An example of channel locations and their corresponding cortical structures are shown in Figure 1.

**Table 1.**
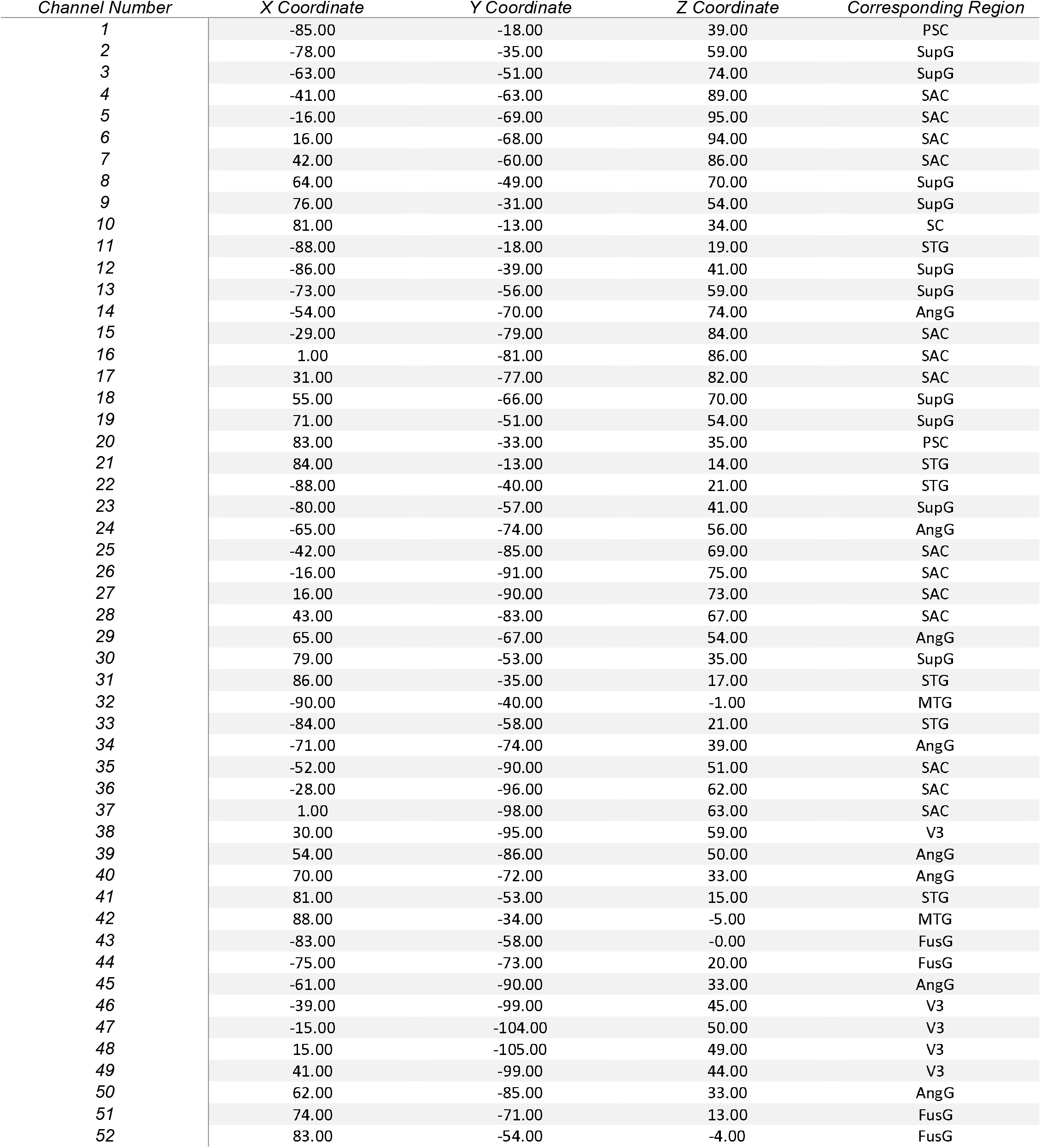
Channel numbers with their corresponding coordinates and regions. Primary somatosensory cortex (PSC), Superior Parietal Gyrus (SupG), Superior Temporal Gyrus (STG), Middle Temporal Gyrus (MTG), Angular Gyrus (AngG), Somatosensory Association Cortex (SAC) and Third Visual Cortex (V3), Fusiform Gyrus (FusG) and Subcentral Area (SC)

| *Channel Number* | *X Coordinate* | *Y Coordinate* | *Z Coordinate* | *Corresponding Region* |
| --- | --- | --- | --- | --- |
| 1 | -85.00 | -18.00 | 39.00 | PSC |
| 2 | -78.00 | -35.00 | 59.00 | SupG |
| 3 | -63.00 | -51.00 | 74.00 | SupG |
| 4 | -41.00 | -63.00 | 89.00 | SAC |
| 5 | -16.00 | -69.00 | 95.00 | SAC |
| 6 | 16.00 | -68.00 | 94.00 | SAC |
| 7 | 42.00 | -60.00 | 86.00 | SAC |
| 8 | 64.00 | -49.00 | 70.00 | SupG |
| 9 | 76.00 | -31.00 | 54.00 | SupG |
| 10 | 81.00 | -13.00 | 34.00 | SC |
| 11 | -88.00 | -18.00 | 19.00 | STG |
| 12 | -86.00 | -39.00 | 41.00 | SupG |
| 13 | -73.00 | -56.00 | 59.00 | SupG |
| 14 | -54.00 | -70.00 | 74.00 | AngG |
| 15 | -29.00 | -79.00 | 84.00 | SAC |
| 16 | 1.00 | -81.00 | 86.00 | SAC |
| 17 | 31.00 | -77.00 | 82.00 | SAC |
| 18 | 55.00 | -66.00 | 70.00 | SupG |
| 19 | 71.00 | -51.00 | 54.00 | SupG |
| 20 | 83.00 | -33.00 | 35.00 | PSC |
| 21 | 84.00 | -13.00 | 14.00 | STG |
| 22 | -88.00 | -40.00 | 21.00 | STG |
| 23 | -80.00 | -57.00 | 41.00 | SupG |
| 24 | -65.00 | -74.00 | 56.00 | AngG |
| 25 | -42.00 | -85.00 | 69.00 | SAC |
| 26 | -16.00 | -91.00 | 75.00 | SAC |
| 27 | 16.00 | -90.00 | 73.00 | SAC |
| 28 | 43.00 | -83.00 | 67.00 | SAC |
| 29 | 65.00 | -67.00 | 54.00 | AngG |
| 30 | 79.00 | -53.00 | 35.00 | SupG |
| 31 | 86.00 | -35.00 | 17.00 | STG |
| 32 | -90.00 | -40.00 | -1.00 | MTG |
| 33 | -84.00 | -58.00 | 21.00 | STG |
| 34 | -71.00 | -74.00 | 39.00 | AngG |
| 35 | -52.00 | -90.00 | 51.00 | SAC |
| 36 | -28.00 | -96.00 | 62.00 | SAC |
| 37 | 1.00 | -98.00 | 63.00 | SAC |
| 38 | 30.00 | -95.00 | 59.00 | V3 |
| 39 | 54.00 | -86.00 | 50.00 | AngG |
| 40 | 70.00 | -72.00 | 33.00 | AngG |
| 41 | 81.00 | -53.00 | 15.00 | STG |
| 42 | 88.00 | -34.00 | -5.00 | MTG |
| 43 | -83.00 | -58.00 | -0.00 | FusG |
| 44 | -75.00 | -73.00 | 20.00 | FusG |
| 45 | -61.00 | -90.00 | 33.00 | AngG |
| 46 | -39.00 | -99.00 | 45.00 | V3 |
| 47 | -15.00 | -104.00 | 50.00 | V3 |
| 48 | 15.00 | -105.00 | 49.00 | V3 |
| 49 | 41.00 | -99.00 | 44.00 | V3 |
| 50 | 62.00 | -85.00 | 33.00 | AngG |
| 51 | 74.00 | -71.00 | 13.00 | FusG |
| 52 | 83.00 | -54.00 | -4.00 | FusG |

**Figure 1.**
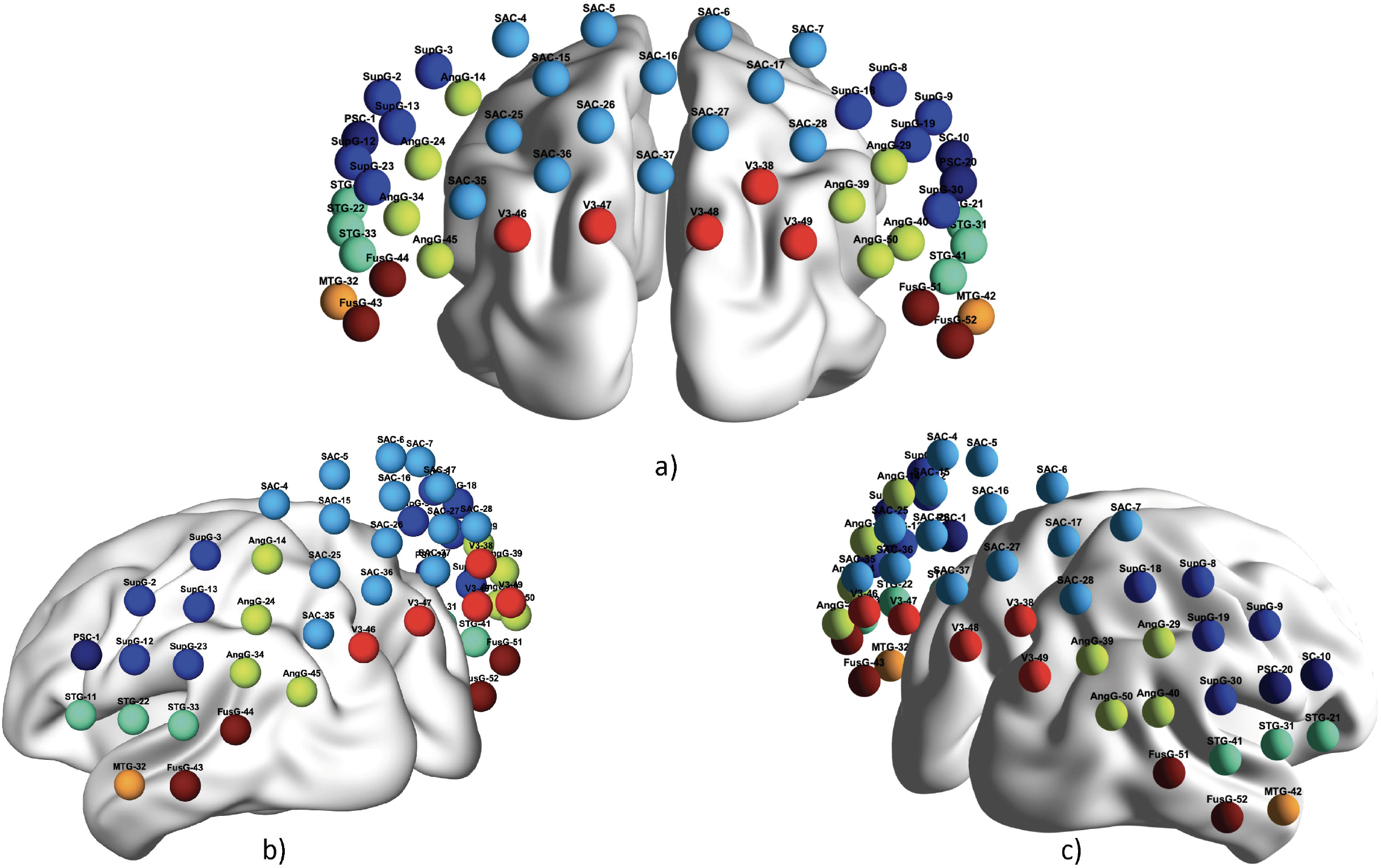
Channel configuration and corresponding cortical structure. For every region, its corresponding channel number was shown after its label. a) Coronal view, b) left hemisplere view c) right hemisphere view. SupG: Supramarginal gyrus, STG: Superior Temporal Gyrus, AngG : Angular Gyrus, MTG : Middle Temporal Gyrus, V3: Third visual cortex, SAC : Somatosensory association cortex, FusG : Fusiform Gyrus.

### 2.3. Data Analysis

Our data preprocessing pipeline includes; band-pass filtering, independent component analysis using fastICA (Hyvarinen, 1999), motion artifact correction using wavelet based-filtering (Molavi & Dumont, 2012) and wavelet-based linear detrending using minimum description length (Jang et al., 2009). We used a 2^nd^ order butterworth band-pass filter with cut-off frequencies between 0.01-0.08 Hz to filter out the heart beat (>1 Hz), respiration (0.15-0.4 Hz) (Fekete et al., 2011) and Mayer waves (∼0.1 Hz) (Yucel et al., 2016) effects. Previous studies revealed that higher correlation values between homologous cortical regions were shown in a wide frequency band (see review (Niu & He, 2014)). After visually inspecting all channels, we noticed some channels were distorted and to overcome this problem, channel interpolation with surrounding channels was performed. To remove motion artifacts, we used wavelet-based motion artifact correction using Daubechies 5 wavelet. Hemodynamic response might include very low frequency artifacts that may influence the functional connections (Di et al., 2013). To remove this effect, we used the wavelet-based-linear detrending using minimum description length algorithm. Independent component analysis was used to separate the neuronal sources from non-neuronal sources.

After performing the preprocessing pipeline, to estimate functional connectivity matrices, we used both CC and Dynamic Time Warping (DTW) distance between all channels for all participants by utilizing ΔHb and ΔHbO time series. In contrast to conventional approach that uses ΔHbO for FC analysis, we also used ΔHb due to previous evidence that suggests both hemoglobin concentrations should be analyzed for sparse networks (Montero-Hernandez et al., 2018b). In addition to this, there are several fNIRS based ML studies that focuses on classification of several psychiatric disorders also uses ΔHb (Cheng et al., 2019; Crippa et al., 2017; Hernandez-Meza et al., 2018; Hernandez-Meza et al., 2017; J. Li et al., 2016; Rosas-Romero et al., 2019; Sirpal et al., 2019; Sutoko et al., 2019) and for some cases ΔHb based features might also show higher accuracy results compared to ΔHbO based ones (Crippa et al., 2017). After estimating the CC based FC matrix, it was normalized by performing Fisher’s z-transformation due to reduce skewness. All preprocessing steps until machine learning process were done using MATLAB (The Mathworks Inc. Natick, MA, USA).

#### 2.3.1. Dynamic Time Warping (DTW)

The other approach, DTW has previously been used in both fMRI (Linke et al., 2020; Meszlenyi, Hermann, et al., 2017), EEG (Karamzadeh et al., 2013) and fNIRS based connectivity studies (Gokcay et al., 2019). DTW is a template matching method used for similarity measurement of two time-varying time series (Sakoe & Chiba, 1978). It is widely used in speech recognition, bioinformatics. It performs an optimum sequence alignment between two time series and it allows to detect identical shapes between time series that may vary in time by performing elastic transformation. For instance, let *S*_1_*=* [*s*_1,0_, *s*_1,1_, *s*_1,2_ ……. *s*_1,*N*− 1,_] and *S*_2_ *=*[*s*_2,0_, *s*_2,1_, *s*_2,2_ ……. *s*_2,*N*− 1,_]be two time series that have the same length *N* and *M* is a distance matrix with *N x N* dimensions. For every index of *M*, we estimate the Euclidean distance between *s*_1,*i*_ and *s*_2,*j*_ as stated Eq. 1;

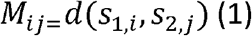

After estimating *M*_*ij*_ that represent the distance for every data point, DTW algorithm tries to find the shortest path between two time series by searching from the index *(0,0)* to *(N* − 1, *N* − 1). The shortest path is called Warping path and represented as *W*.

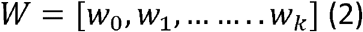

*W* is constructed by using linear programming. First, euclidean distance *d(s*_1,*i*_, *s*_2,*j*_ ) should be found and defined as cost value to estimate the minimum distance between two time series. To move forward in the warping matrix, minimum value of the neighboring cells (*M(i* − 1, *j* − 1*), M(i* − 1, *j), M(i, j* − 1))) is selected and the *M(i, j)* is calculated by summing of *d(s*_1_,_*i*_ *s*_2,*j*_) and minimum value as shown in Eq (3).

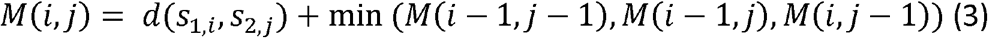

If we define indices of the warping matrix as *p* and *r* for one time series and *x* and *y* for the other one, *t*^th^ and *t* − 1^th^ point in our warping path can be identified as *w = (p, r)* and *w*_t − 1_*= (p′, r′)* and this warping path must satisfy following four constraints;

Monotonicity constraint: In warping path indices either increase or stay same in time domain. Therefore, it will not be possible to repeat the warping path. Indices ensure the following condition as shown in Eq. (4)

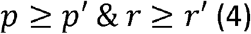

Continuity constraint: Warping path advances one step at a time. Index change between *p* - *p′* and *r* - *r′* can be less or equal to one. Equation 5 shows this condition.

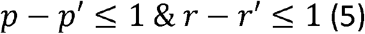

Boundary constraint: The path initiates from value *M*(0,0) and ends in value *M*(*N* − 1, *N* − 1). Warping window constraint: An intuitive alignment path onto distance matrix *M* is not expected to drift from the diagonal. For warping window length *l*,

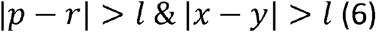

We chose warping window length *l* as 1 sec (10 samples). After estimating the DTW distance for functional connectivity, as it was previously stated in (Linke et al., 2020; Meszlenyi, Hermann, et al., 2017), we first normalized the distance by dividing it to length of the time series. Then, we multiplied the values to -1 and demeaned to a transform distance. If this transformed distance is greater than 0, it is represented as above average similarity and if it is lower than 0, it is represented as below average similarity.

#### 2.3.2. Machine Learning

First, we separately used the extracted features from ΔHb and ΔHbO and performed classification. In this study, we defined the correlation coefficient based ΔHb and ΔHbO feature set as CC-ΔHb and CC-ΔHbO. For DTW, we defined the feature sets as DTW-ΔHb and DTW-ΔHbO. Then, we concatenated the feature sets for both ΔHb and ΔHbO and performed the same procedure again for both connectivity measures as DTW-Fuse and CC-Fuse. All machine learning steps including, feature selection, parameter optimization and classification were carried out by using Python based Scikit-Learn toolkit (Pedregosa et al., 2011).

##### 2.3.2.1. Feature Selection

Before feature selection, we first converted the 52 x 52 connectivity matrix to 52 x 51 /2 = 1326 pairwise connectivity features. Every connection was considered as a feature. Therefore, for every dataset (CC-ΔHbO, CC-ΔHb, DTW-ΔHbO and DTW-ΔHb), we had a 43 x 1326 feature vector. To select optimum features from every dataset, we used L1-norm Support Vector Machine (SVM). For L1-norm SVM based feature selection, the regularization parameter (C) controls the sparsity and as we increase C value, we get more features. For optimum number of features to avoid overfitting, we used C = 0.1.

For classification of flourishing individuals, we selected connections for CC-ΔHbO as L SMG – L SMG, R V3 – R SAC, R STG – L STG, L AngG – L SAC, for CC-ΔHb as L STG – L MTG, L FusG – L MTG, L FusG – L AngG. For DTW measure, we selected connections for DTW-ΔHbO as L SAC – R SMG, RAngG – R SCA, L FusG – R SMG, L FusG – R SAC, R FusG – R SMG and for DTW-ΔHb as L SAC-L SAC, L FusG - L SCA, L FusG – R SMG, L V3 – L SAC. After feature selection step, we also created a third feature vector for every connectivity measure for classification of flourishing individuals by fusing the ΔHbO and ΔHb results. For CC, we created CC-Fuse by concatenating the CC-ΔHbO with CC-ΔHb and for DTW, we created DTW-Fuse by concatenating DTW-ΔHbO with DTW-ΔHb.

##### 2.3.2.2. Hyperparameter-Optimized Classification using Nested Cross-Validation

After selecting features, we performed nested cross-validation that includes hyperparameter optimization as inner loop and classifier generalization step in outer loop. After splitting the dataset as training and test dataset, in the inner loop, we trained the model using training dataset and found the optimal hyperparameters for the classifier. In the outer loop, we built the model with the optimal hyperparameters from the inner loop and tested it by using the test dataset. We did not include the feature selection into this nested cross validation due to having low number of samples (N=43) and to observe whether our selected features can be potential biomarkers that discriminates highly/normal flourishing individuals or not. We used 10 fold for inner loop and 10 fold for outer loop in nested cross-validation for 30 times and averaged the scores. Whole data analysis pipeline is shown in Figure 2.

**Figure 2.**
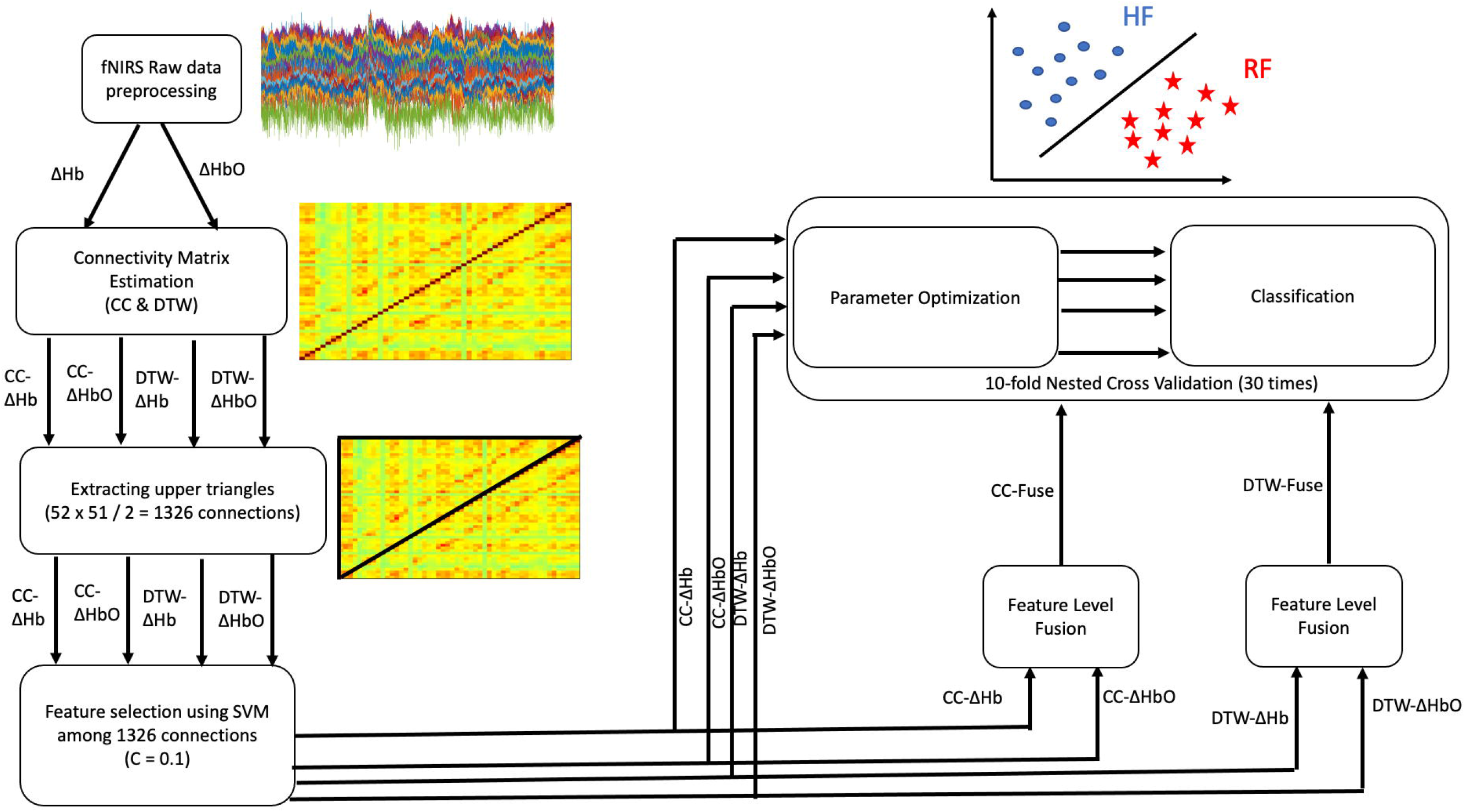
Data Analysis Flow Chart. CC - ΔHbO: Feature set of ΔHbO using Pearson’s correlation, CC - ΔHb : Feature set of ΔHb using Pearson’s correlation, CC - Fuse: Feature set of fusion of ΔHb and ΔHbO feature sets using Pearson’s correlation, DTW - ΔHbO : Feature set of ΔHbO using Dynamic Time Warping, DTW - ΔHb : Feature set of ΔHb using Dynamic Time Warping, DTW - ΔHbO : Feature set of ΔHbO using Dynamic Time Warping, DTW - Fuse: Feature set of fusion of ΔHb and ΔHbO feature sets using Dynamic Time Warping.

We used several classifiers such as SVM with Linear and Radial Basis Function (RBF) kernels, Linear Discriminant Analysis (LDA), Quadratic Discriminant Analysis (QDA), k-nearest neighborhood, Gradient Boosting, AdaBoost, Naïve Bayes and Logistic Regression. For every algorithm, we used several parameters to optimize the classifier. All those parameters were reported in Table 2. To find the optimal parameters of classifiers, we used grid-search algorithm by using GridSearchCV() function in scikit-learn toolkit to carry out this operation. We also extracted receiver operating characteristic (ROC) curves and estimated Area Under Curve (AUC) values for every feature set and every classifier. We accepted the AUC greater than 0.9 as success rate due to being accepted as excellent (0.9-1) according to the previous studies (El Khouli et al., 2009; Ludemann et al., 2006; Metz, 1978; Obuchowski, 2003).

**Table 2.** Used hyperparameters for classifiers.

| *Classifier* | *Parameter* |
| --- | --- |
| *Nearest neighbor* | • Nearest neighbor : 2,3...,10<br>• Weights : Uniform, distance |
| *Linear SVM* | • C : 0.001, 0.01, 0.1, 1, 10, 100<br>• Gamma: 0.001, 0.01, 0.1, 1<br>• Class weight : balanced, none |
| *RBF SVM* | • C : 0.001, 0.01, 0.1, 1, 10, 100<br>• Gamma: 0.001, 0.01, 0.1, 1<br>• Class weight : balanced, none |
| *Gradient Boosting* | • Number of estimators : 50, 100<br>• Learning rate : 0.001, 0.005, 0.01, 0.01, 0.05 |
| *AdaBoost* | • Number of estimators : 50, 100<br>• Learning rate : 0.001, 0.005, 0.01, 0.01, 0.05 |
| *Naïve Bayes* | No parameter |
| *Linear Discriminant Analysis* | No parameter |
| *Quadratic Discriminant Analysis* | No parameter |
| *Logistic Regression* | • Penalty : L1 and L2 regularization<br>• Alpha : 0.01, 0.1<br>• Number of iteration : 1,2,5,10,100,500 |

##### 2.3.2.3. Statistical Analysis

After performing the classification process, Pearson’s correlation was carried out between all features of four flourishing feature sets (CC-ΔHbO, CC-ΔHb, DTW-ΔHbO, DTW-ΔHb) and clinical data. For statistical significance, we defined p value as 0.001.

## 3. Results

### 3.1. Classification Accuracy Results

For CC based features, CC-Fuse showed higher accuracy scores for all classifiers except for AdaBoost classifier compared to CC-ΔHbO and CC-ΔHb. Among all classifiers, highest accuracy was found by using Linear SVM and LDA for CC-Fuse (mean ± std: 0.89 ± 0.01). Among CC-ΔHb and CC-ΔHbO classification results, we found the highest accuracy by using LDA and logistic regression (mean ± std: 0.81 ± 0.01). Except for gradient boosting classifier, CC-ΔHbO feature set yield higher accuracy results compared to CC-ΔHb feature set.

For DTW based features, we found the highest accuracy by using DTW-Fuse dataset and Naïve Bayes classifier (mean ± std: 0.89 ± 0.01). Among DTW-ΔHbO and DTW-ΔHb classification results, we found the highest accuracy by using DTW-ΔHb feature set and LDA classifier (mean ± std: 0.82 ± 0.01). When we compared the DTW-ΔHbO and DTW-ΔHb classification results, DTW-ΔHb based classification accuracies are higher than DTW-ΔHbO based ones for all classifiers. All accuracy results of flourishing individuals are shown in Table 3 and all accuracy violin plots were shown in Figure 3.

**Table 3.**
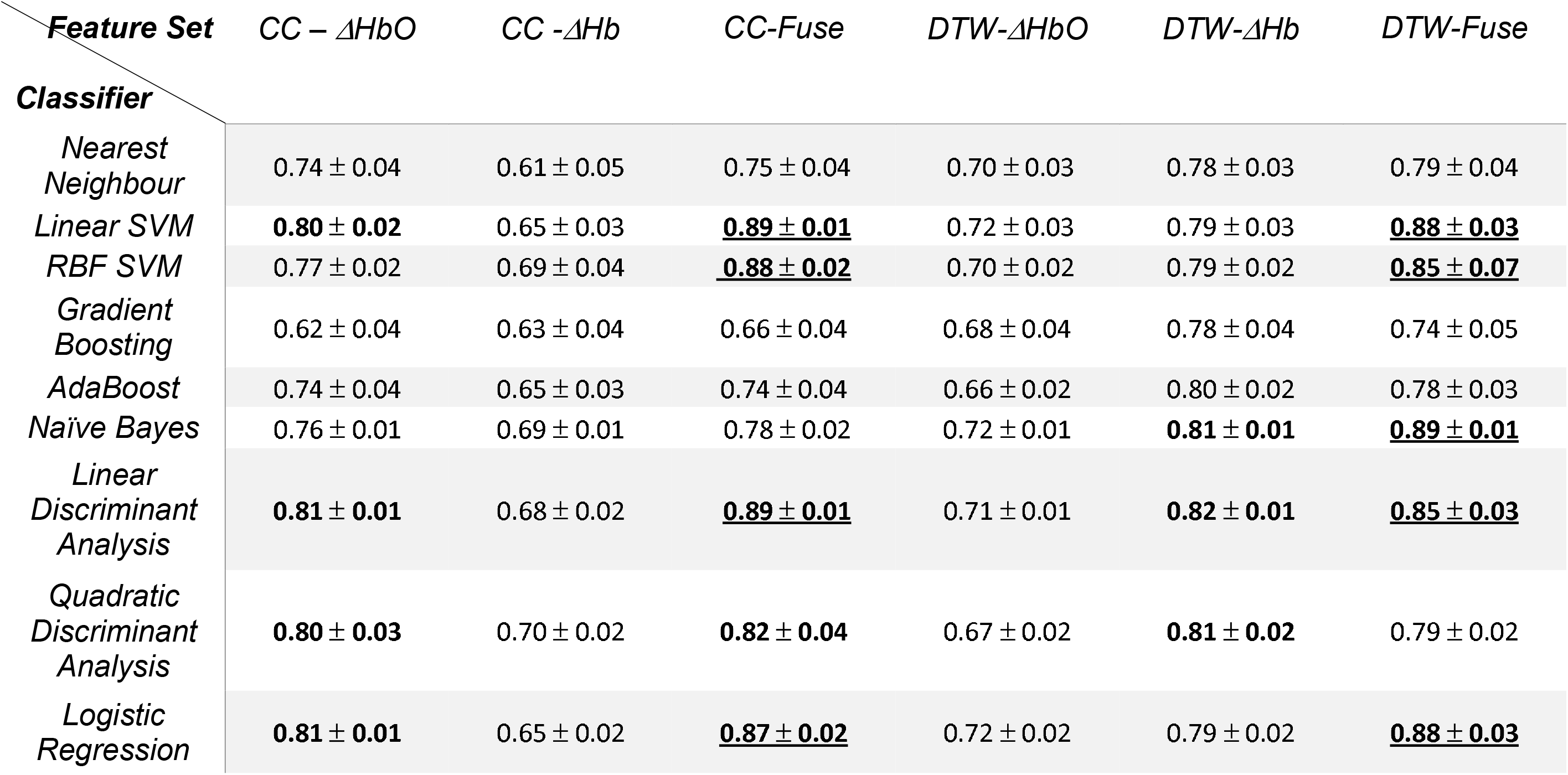
Mean and Standard deviation accuracy results for classification of highly flourishing and normal flourishing participants using different classifiers. CC: Correlation coefficient, ΔHb : Deoxyhemoglobin concentration change, ΔHbO : Oxyhemoglobin concentration change. SVM : Support Vector Machine, RBF : Radial Basis Function, DTW : Dynamic Time Warping. Bold highlighted results are the accuracy values greater than 80 % and bold highlighted and underlined results are the accuracy values greater than 85 %.

| **Feature Set** | **CC – $\Delta HbO$** | **CC -$\Delta Hb$** | **CC-Fuse** | **DTW-$\Delta HbO$** | **DTW-$\Delta Hb$** | **DTW-Fuse** |
| --- | --- | --- | --- | --- | --- | --- |
| **Classifier** |  |  |  |  |  |  |
| *Nearest Neighbour* | 0.74 ± 0.04 | 0.61 ± 0.05 | 0.75 ± 0.04 | 0.70 ± 0.03 | 0.78 ± 0.03 | 0.79 ± 0.04 |
| *Linear SVM* | **0.80 ± 0.02** | 0.65 ± 0.03 | **<u>0.89 ± 0.01</u>** | 0.72 ± 0.03 | 0.79 ± 0.03 | **<u>0.88 ± 0.03</u>** |
| *RBF SVM* | 0.77 ± 0.02 | 0.69 ± 0.04 | **<u>0.88 ± 0.02</u>** | 0.70 ± 0.02 | 0.79 ± 0.02 | **<u>0.85 ± 0.07</u>** |
| *Gradient Boosting* | 0.62 ± 0.04 | 0.63 ± 0.04 | 0.66 ± 0.04 | 0.68 ± 0.04 | 0.78 ± 0.04 | 0.74 ± 0.05 |
| *AdaBoost* | 0.74 ± 0.04 | 0.65 ± 0.03 | 0.74 ± 0.04 | 0.66 ± 0.02 | 0.80 ± 0.02 | 0.78 ± 0.03 |
| *Naïve Bayes* | 0.76 ± 0.01 | 0.69 ± 0.01 | 0.78 ± 0.02 | 0.72 ± 0.01 | **0.81 ± 0.01** | **<u>0.89 ± 0.01</u>** |
| *Linear Discriminant Analysis* | **0.81 ± 0.01** | 0.68 ± 0.02 | **<u>0.89 ± 0.01</u>** | 0.71 ± 0.01 | **0.82 ± 0.01** | **<u>0.85 ± 0.03</u>** |
| *Quadratic Discriminant Analysis* | **0.80 ± 0.03** | 0.70 ± 0.02 | **0.82 ± 0.04** | 0.67 ± 0.02 | **0.81 ± 0.02** | 0.79 ± 0.02 |
| *Logistic Regression* | **0.81 ± 0.01** | 0.65 ± 0.02 | **<u>0.87 ± 0.02</u>** | 0.72 ± 0.02 | 0.79 ± 0.02 | **<u>0.88 ± 0.03</u>** |

**Figure 3.**
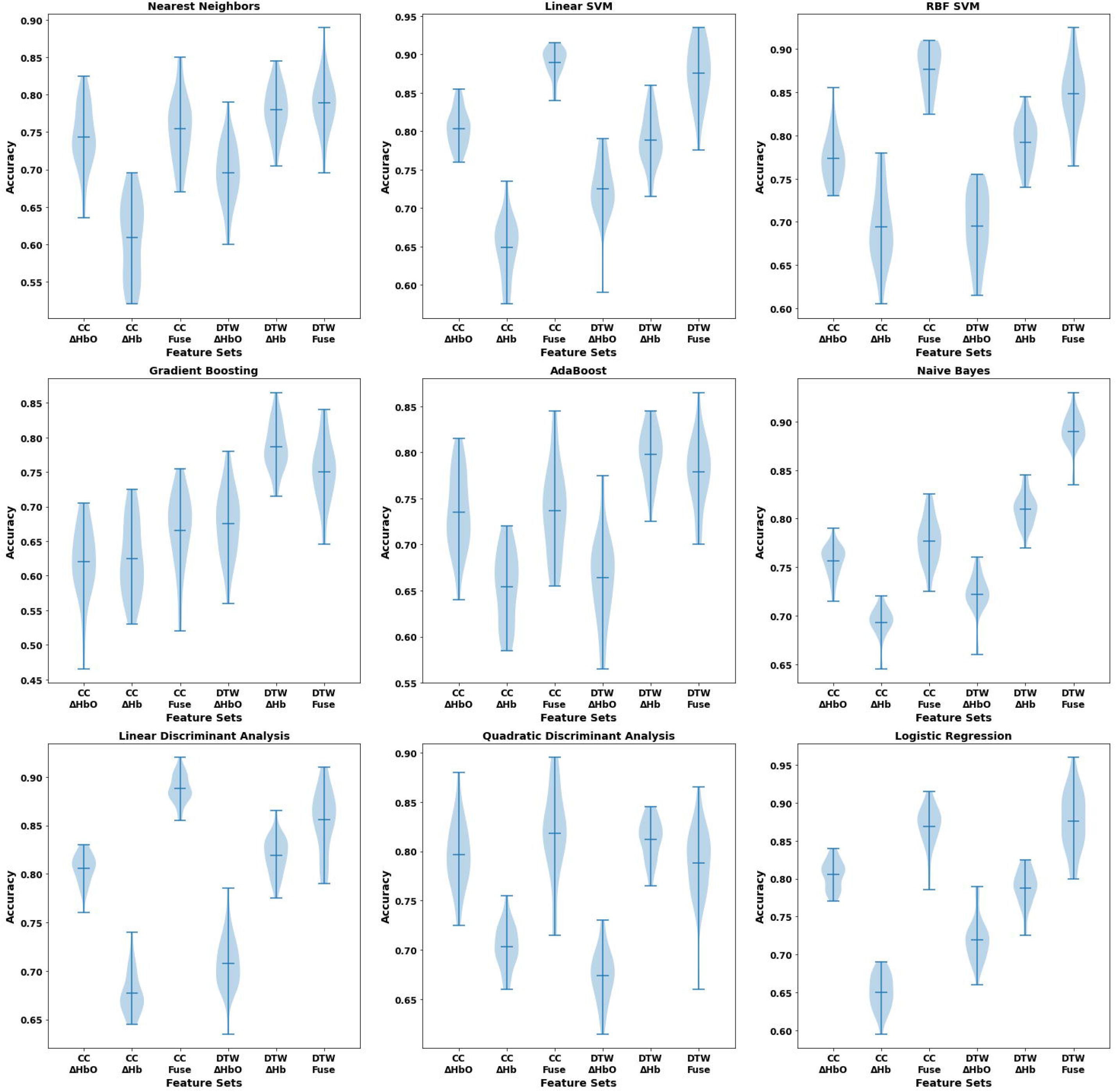
Accuracy violin plots for every classifier. CC - ΔHbO: Feature set of ΔHbO using Pearson’s correlation, CC - ΔHb : Feature set of ΔHb using Pearson’s correlation, CC - Fuse: Feature set of fusion of ΔHb and ΔHbO feature sets using Pearson’s correlation, DTW - ΔHbO : Feature set of ΔHbO using Dynamic Time Warping, DTW - ΔHb : Feature set of ΔHb using Dynamic Time Warping, DTW - ΔHbO : Feature set of ΔHbO using Dynamic Time Warping, DTW - Fuse: Feature set of fusion of ΔHb and ΔHbO feature sets using Dynamic Time Warping. RBF SVM : Radial Basis Function Support Vector Machine.

AUC results for classification of flourishing individuals showed that there are several classifiers and feature sets that showed AUC score more than 0.9. Among all classifiers and feature sets, highest AUC value was found by using DTW-Fuse feature set and Naïve Bayes classifier as 0.95. AUC results of all classifiers and the corresponding ROC curves are shown in Table 4 and Figure 4 respectively.

**Table 4.** Area under curve (AUC) results for classification of highly flourishing and normal flourishing participants using different classifiers. CC: Correlation coefficient, ΔHb : Deoxyhemoglobin concentration change, ΔHbO : Oxyhemoglobin concentration change. SVM : Support Vector Machine, RBF : Radial Basis Function, DTW : Dynamic Time Warping. Bold highlighted and underlined results are the AUC values greater than 0.9.

| **Feature Set** | **CC – ΔHbO** | **CC -ΔHb** | **CC-Fuse** | **DTW-ΔHbO** | **DTW-ΔHb** | **DTW-Fuse** |
| --- | --- | --- | --- | --- | --- | --- |
| **Classifier** |  |  |  |  |  |  |
| *Nearest Neighbour* | 0.82 | 0.68 | 0.84 | 0.73 | 0.83 | 0.84 |
| *Linear SVM* | 0.82 | 0.66 | **<u>0.90</u>** | 0.79 | 0.86 | **<u>0.91</u>** |
| *RBF SVM* | 0.83 | 0.69 | **<u>0.92</u>** | 0.75 | 0.87 | **<u>0.90</u>** |
| *Gradient Boosting* | 0.65 | 0.69 | 0.66 | 0.74 | 0.81 | 0.77 |
| *AdaBoost* | 0.76 | 0.68 | 0.81 | 0.66 | 0.87 | 0.82 |
| *Naïve Bayes* | 0.81 | 0.74 | 0.86 | 0.82 | **<u>0.93</u>** | **<u>0.95</u>** |
| *Linear Discriminant Analysis* | 0.86 | 0.72 | **<u>0.91</u>** | 0.79 | 0.88 | 0.89 |
| *Quadratic Discriminant Analysis* | 0.87 | 0.75 | 0.87 | 0.77 | **<u>0.92</u>** | 0.86 |
| *Logistic Regression* | 0.87 | 0.71 | **<u>0.90</u>** | 0.78 | 0.86 | **<u>0.91</u>** |

**Figure 4.**
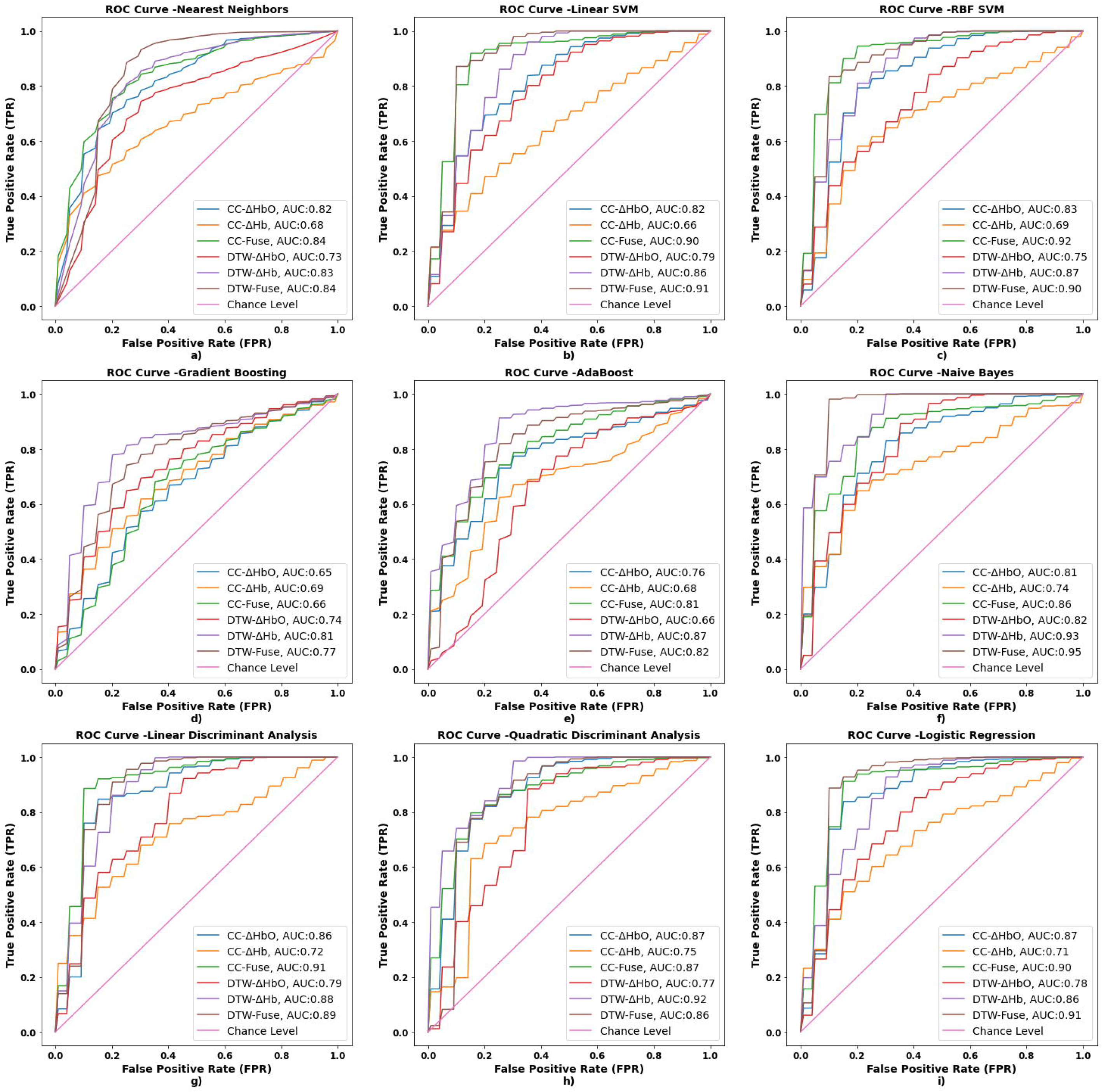
Receiver operating characteristic (ROC) curves for every classifier and every feature set. CC - ΔHbO: Feature set of ΔHbO using Pearson’s correlation, CC - ΔHb : Feature set of ΔHb using Pearson’s correlation, CC - Fuse: Feature set of fusion of ΔHb and ΔHbO feature sets using Pearson’s correlation, DTW - ΔHbO : Feature set of ΔHbO using Dynamic Time Warping, DTW - ΔHb : Feature set of ΔHb using Dynamic Time Warping, DTW - ΔHbO : Feature set of ΔHbO using Dynamic Time Warping, DTW - Fuse: Feature set of fusion of ΔHb and ΔHbO feature sets using Dynamic Time Warping. RBF SVM : Radial Basis Function Support Vector Machine. AUC : Area under curve.

#### 3.1.1. Classifying Flourishing Individuals by Using Both Connectivity Measures

Having noticed the efficiency of CC-ΔHbO and DTW-ΔHb based features for classification of flourishing individuals, we created another type of feature set by fusing these two feature sets. We performed classification by following the same paradigm mentioned above and we found the highest accuracy as 90% by using Nearest Neighbour and RBF SVM with 0.90 and 0.93 AUC. All accuracy values are numerically shown in Table 5 and also are shown as violin plots in Figure 5. Also, corresponding ROC curves were shown in Figure 6.

**Table 5.** Mean and Standard deviation accuracy results and Area Under Curve (AUC) results for classification of highly flourishing and normal flourishing participants using different classifiers and fused features from different connectivity metrics. AUC: Area Under Curve, CC: Correlation coefficient, ΔHb : Deoxyhemoglobin concentration change, ΔHbO : Oxyhemoglobin concentration change. SVM : Support Vector Machine, RBF : Radial Basis Function, DTW : Dynamic Time Warping. Bold highlighted results are the accuracy values greater than 80 % and bold highlighted and underlined results are the accuracy values greater than 85 %.

| *Feature Set*<br>*Classifier* | *CC – ΔHbO &*<br>*DTW-ΔHb*<br>*Accuracy* | *CC – ΔHbO*<br>*&*<br>*DTW-ΔHb*<br>*AUC* |
| --- | --- | --- |
| *Nearest Neighbor* | **<u>0.90 ± 0.01</u>** | 0.90 |
| *Linear SVM* | **<u>0.87 ± 0.02</u>** | 0.91 |
| *RBF SVM* | **<u>0.90 ± 0.03</u>** | 0.94 |
| *Gradient Boosting* | 0.74 ± 0.03 | 0.78 |
| *AdaBoost* | 0.79 ± 0.03 | 0.88 |
| *Naïve Bayes* | **<u>0.85 ± 0.02</u>** | 0.95 |
| *Linear Discriminant*<br>*Analysis* | **<u>0.85 ± 0.01</u>** | 0.93 |
| *Quadratic*<br>*Discriminant*<br>*Analysis* | **<u>0.85 ± 0.02</u>** | 0.92 |
| *Logistic Regression* | **<u>0.86 ± 0.01</u>** | 0.90 |

**Figure 5.**
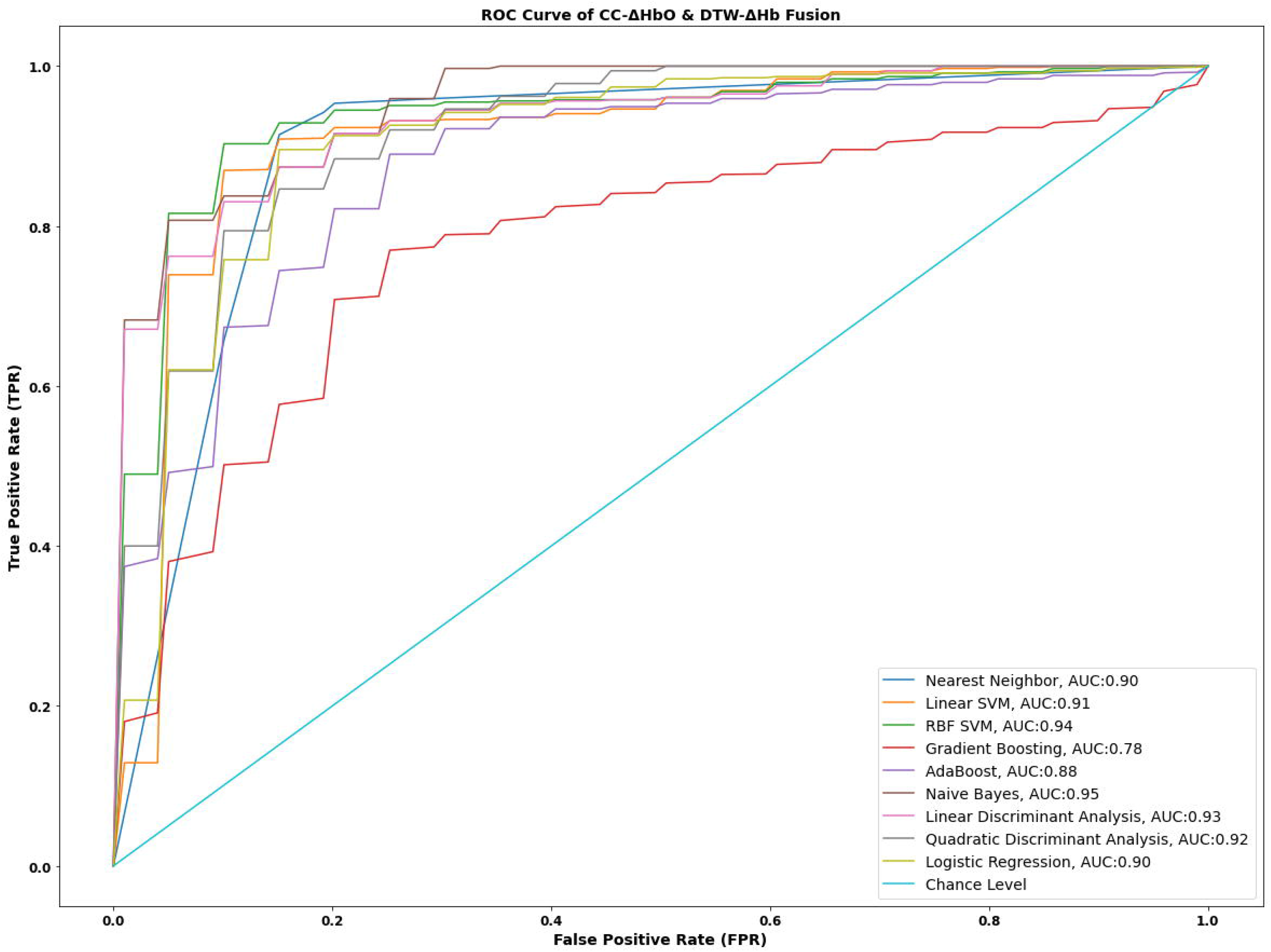
Receiver operating characteristic (ROC) curves for every classifier for fusion of CC - ΔHbO and DTW - ΔHb feature sets. AUC : Area under curve

**Figure 6.**
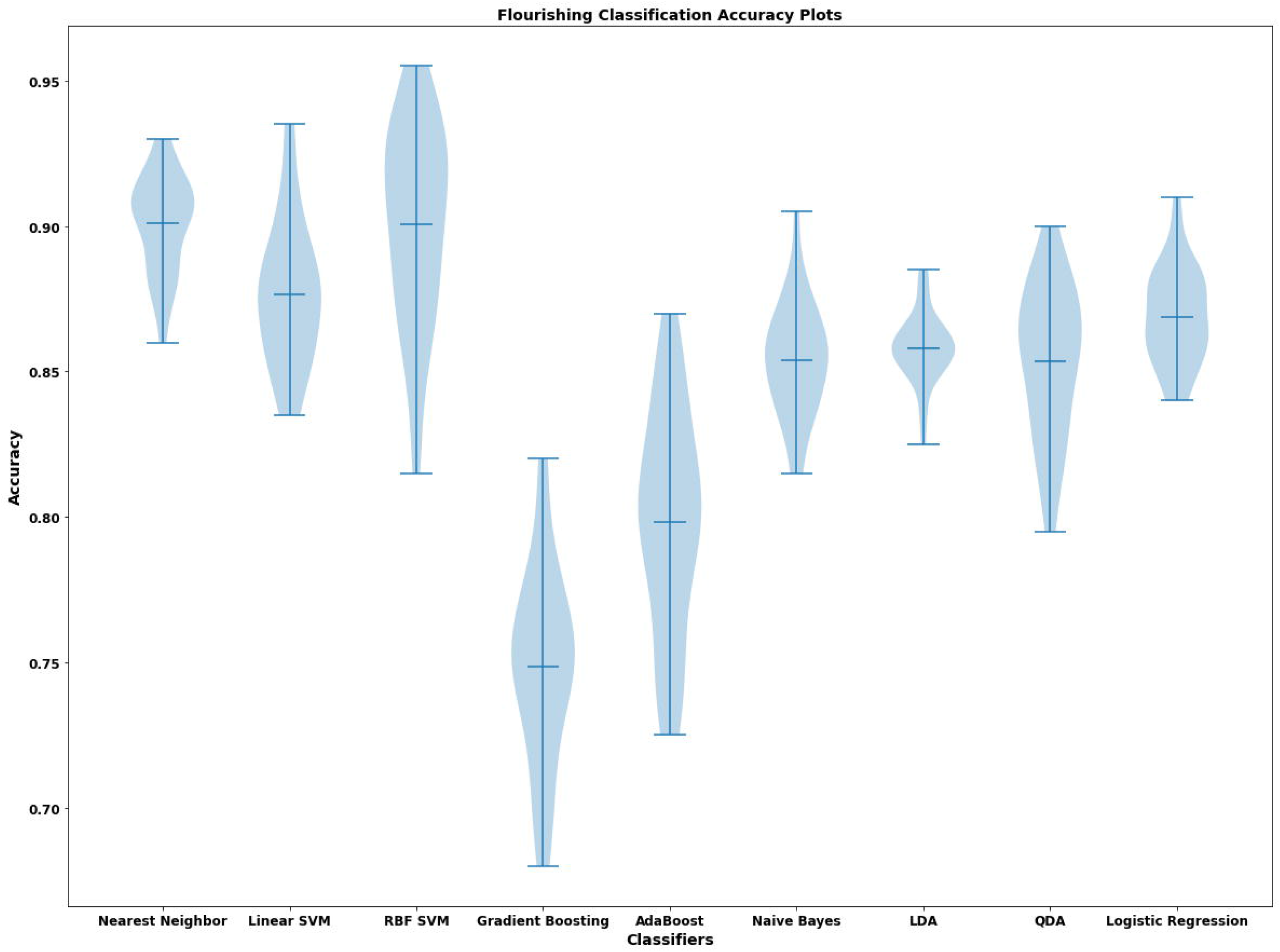
Accuracy violin plots for every classifier for fusion of CC - ΔHbO and DTW - ΔHb feature sets. LDA : Linear Discriminant Analysis, QDA : Quadratic Discriminant Analysis.

### 3.2. Association Between Features and Clinical Variables

For all clinical data, we both analyzed the correlation between CC and DTW based features. We found negative significant correlation between flourishing score and CC-ΔHbO measure of R V3 – R SAC connection (r = -0.551, p < 0.001) and DTW-ΔHb measure of L V3 – L SAC connection (r = -0.496, p < 0.001). Correlation plots between connection measures and flourishing scores are shown in Figure 7.

**Figure 7.**
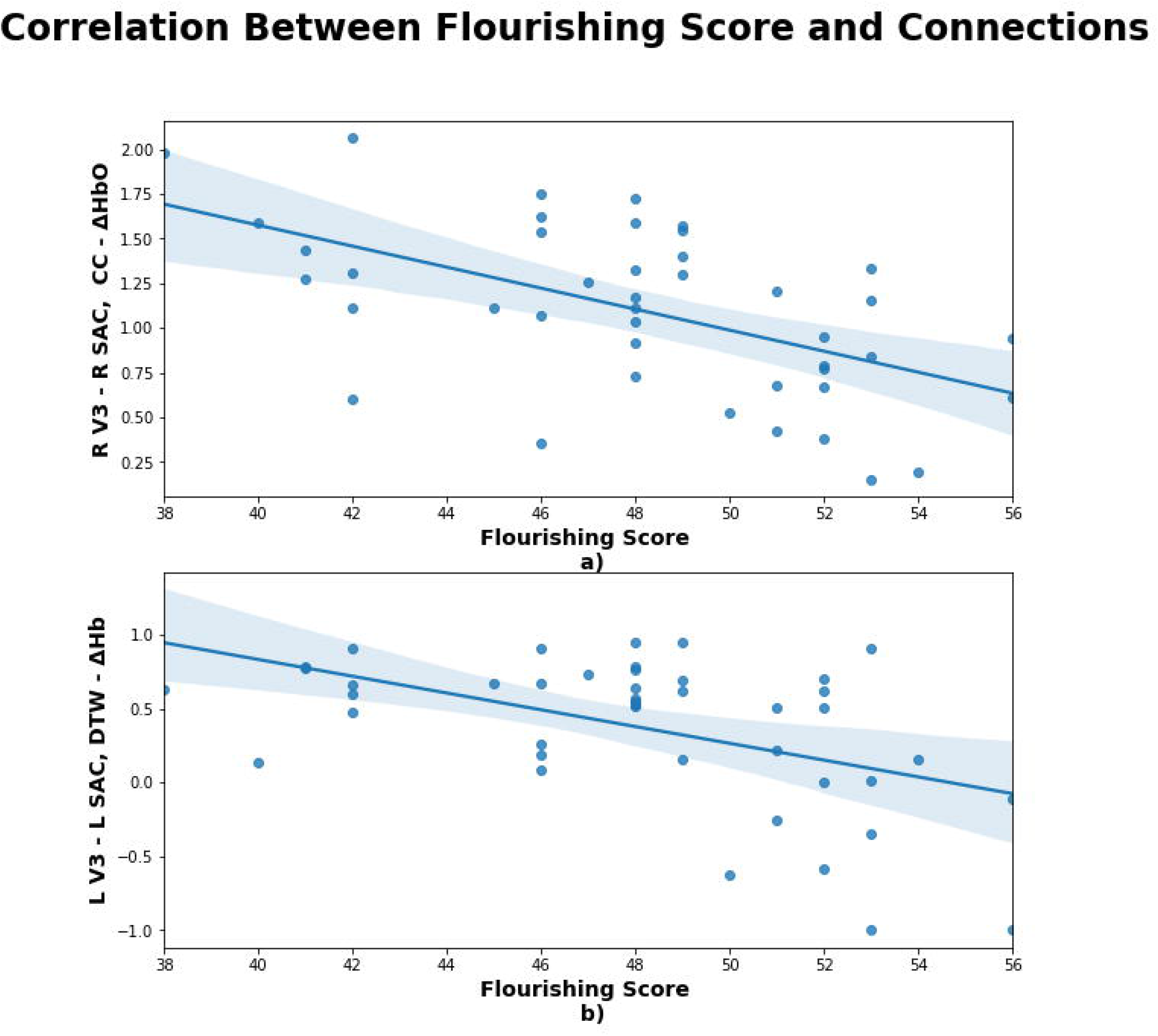
Correlation plots between flourishing scores and R V3 – R SAC using CC - ΔHbO (a) and L V3 – L SAC using DTW - ΔHb (b). In a) the CC values were Z-transformed.

## 4. Discussion

In this study, we investigated the neural correlates of flourishing levels of individuals by using a previously collected resting-state fNIRS data and different machine learning approaches. We classified both groups (HF / NF) with high accuracy and AUC values. In addition to conventional connectivity measure CC, we utilized DTW algorithm to try to extract discriminative features. We classified individuals with 90% accuracy by fusing the CC-ΔHbO and DTW-ΔHb based connections. To our best knowledge, our study includes several novelties such that being the first study that neuroimaging data was used to classify flourishing individuals via ML techniques and being the first study that compares efficiency of DTW and CC using ΔHb and ΔHbO signals on classification of a resting state fNIRS data.

### 4.1. Efficiency of Connectivity Measures

We utilized both DTW and CC measures to estimate functional connectivity in this study and found that DTW-ΔHb dataset gave higher accuracy than DTW-ΔHbO and CC-ΔHbO dataset gave higher accuracy than CC-ΔHb dataset for classification of flourishing. Also, 7 of 9 classifiers gave higher accuracy results for DTW-ΔHb dataset than for CC-ΔHbO dataset. Compared to CC, DTW is a novel measure in functional connectivity estimation and few studies were performed by utilizing DTW for EEG (Karamzadeh et al., 2013), fMRI (Jin et al., 2020; Linke et al., 2020; Meszlenyi, Hermann, et al., 2017; Mohanty et al., 2020) and fNIRS (Gokcay et al., 2019). In a recent study, it was shown that its efficiency outperformed conventional CC approach to detect atypical connectivity patterns in autism and it was reported that DTW was found sensitive to BOLD signal amplitude which is dependent on Hb concentration (Linke et al., 2020). In another study that uses DTW and CC to classify ADHD patients and gender groups, it was reported that DTW showed better performance for classification using SVM and Least Absolute Shrinkage and Selection Operator (LASSO) (Meszlényi et al., 2016). Similar results were also found for classification of amnestic mild cognitive impairment patients using rsfMRI and Convolutional Neural Network (Meszlenyi, Buza, et al., 2017). Also, it was reported that DTW showed more stable FC than conventional CC and also captured the non-stationarity in simulated fMRI data (Meszlenyi, Hermann, et al., 2017). Unlike CC, DTW allows to catch similar but delayed changes between time series, therefore it can detect the non-linear behaviors of time series (Karamzadeh et al., 2013). On the other hand, in case of such a lag, CC will be low regardless of which time series it was used and this might directly lower the classification accuracy.

We also found that DTW-ΔHb and CC-ΔHbO feature sets showed higher classification scores for classification of flourishing levels of individuals. In general, fNIRS based connectivity studies generally utilizes ΔHbO and ΔHbO signals have higher signal-to-noise ratio (SNR) than ΔHb signals (Homae et al., 2010; Montero-Hernandez et al., 2018a; Niu et al., 2011; Y. J. Zhang et al., 2010). On the other hand, having low SNR of ΔHb signals might cause this difference due to less affection from extracerebral and intra cerebral systemic artifacts (Kirilina et al., 2012). Despite performing several preprocessing steps, short-channel regression could not be performed to remove extracerebral systemic artifacts out. Another finding was to obtain highest accuracy result by combining the feature sets DTW-ΔHb and CC-ΔHbO. Our result supports the idea to combine the feature sets extracted from both connectivity measures for classification to obtain higher accuracy compared to using one type of connectivity measure (Meszlenyi, Buza, et al., 2017). Both measures have their own pros and cons and they should be used together to reveal the potential differences between groups due to focusing different aspects of functional networks (Linke et al., 2020). Further basic research is needed to reveal the reason that causes this difference.

### 4.2. Selected Features as Potential Biomarkers

After extracting functional connections between regions, we found discriminative regions for classification of flourishing levels. We utilized connections L SMG – L SMG, R V3 – R SAC, R STG – L STG, L AngG – L SAC for CC-ΔHbO, L STG – L MTG, L FusG – L MTG, L FusG – L AngG for CC-ΔHb, L SAC – R SMG, RAngG – R SCA, L FusG – R SMG, L FusG – R SAC, R FusG – R SMG for DTW-ΔHbO and L SAC-L SAC, L FusG - L SCA, L FusG – R SMG, L V3 – L SAC for DTW-ΔHb. As it was previously stated, these regions are strongly involved in DMN (Greicius et al., 2003; Raichle et al., 2001). Among these regions, right and left STG was strongly associated to social well-being according to a recent study (Kong, Xue, et al., 2016) and STG plays an important role social perception (Kanai et al., 2012; Yang et al., 2015). Also, by using SVM as a feature extraction technique instead of conventional statistical approaches, L MTG – L FusG connection was found as a reproduced result to discriminate flourishing (F. Goldbeck et al., 2018). FusG is widely known about its role in face perception (Grill-Spector et al., 2017). A previous PET study reported that baseline glucose metabolism in bilateral FusG, STG and MTG was found significantly associated to subjective well-being (Volkow et al., 2011).

In parietal region, L SMG, the subpart of left inferior parietal lobe (IPL), was strongly associated with well-being (Luo et al., 2016; Volkow et al., 2011; Waytz et al., 2015). The other subpart of inferior parietal lobe, AngG was found strongly related to subjective happiness (Katsumi et al., 2020). Association between well-being and IPL might be related to the role of IPL in episodic memory (Liang et al., 2012; Wagner et al., 2005). On the other hand, a very recent study related to relationship between rsfMRI and subjective happiness showed that right SAC (BA 7) and amygdala connection were strongly associated with subjective happiness (W. Sato et al., 2019). Like IPL, SAC is also related to episodic memory (see review (Cabeza et al., 2008)). These regions suggested that subcomponents of DMN might be potential neural markers to identify levels of flourishing for individual level.

### 4.3. Classification Results

Among classification results, utilizing both CC-ΔHbO and DTW-ΔHb datasets, we found highest accuracy (90%) with 0.90 and 0.94 AUC values by using nearest neighbor and RBF SVM classifier, respectively. To our best knowledge, there is no study that uses neural data to classify levels of well-being however, there are two rsfMRI studies that uses neural data to predict well-being (Kong, Wang, et al., 2016; Kong, Xue, et al., 2016). In the first study, eudaimonic well-being was predicted by using regional homogeneity in inferior frontal gyrus and using linear regression with 4-fold cross validation (Kong, Wang, et al., 2016). The other study showed that right posterior STG and thalamus was used to predict social well-being using mediation analysis (Kong, Xue, et al., 2016). Also, there are some studies that utilizes behavioral and physiological data to both predict the levels of well-being. A recent study that utilized behavioral data collected via online survey such as Adolescent Self-Rating Life Events Check List, Big Five Inventory, Child and Adolescent Social Support Scale found 90% accuracy with 92% sensitivity and 90% specificity (N. Zhang et al., 2019). Also, a well-being level prediction study that considered a physical, mental and general health index by utilizing domotic sensor data and ML techniques revealed that mean absolute error of prediction of physical, mental and general health index were found as 32%, 13% and 17%, respectively (Casaccia et al., 2020). In another study, physiological data (heart rate, skin conductance), phone, mobility and modifiable behavior features gave 78% accuracy for classification of high or low stress, 86% accuracy for classification of high or low mental health, and 74% accuracy for classification of high or low stress groups (Sano et al., 2018). By utilizing laboratory (blood pressure, cholesterol etc.), demographic (age, gender, race, weight etc.) and lifestyle (frequency of drinking alcohol, vigorous work activity, number of cigarettes per day etc.) data and using machine learning algorithms such as SVM, Bagging, Adaboost, Random Forest etc., highest AUC value was found as 0.726 (Agarwal et al., 2016). Compared to these studies, our results are notable and outperformed some previous studies that was used several different objective measures. Also, while reaching these accuracy values, we utilized neural data which is an objective measure.

On the other hand, compared to fMRI, fNIRS has several advantages such as ease of use, mobility, being inexpensive and there are several studies that utilizes fNIRS data to classify several psychiatric disorders such as depression (Husain et al., 2020; Takizawa et al., 2014; Zhu et al., 2020), schizophrenia (Azechi et al., 2010; Chuang et al., 2014; Dadgostar et al., 2018; Einalou et al., 2016; Hahn et al., 2013; Ji et al., 2020; Koike et al., 2017; Z. Li et al., 2015; Song et al., 2017) using machine learning techniques. High classification accuracies revealed that fNIRS might be a promising tool to identify the levels of flourishing and also closely related to other psychiatric disorders (Baskak, 2018; Ehlis et al., 2014).

### 4.4. Correlation Results Between Flourishing Scores and Used Features

We found negative significant correlations between flourishing score and CC-ΔHbO measure of R V3 – R SAC connection and DTW-ΔHb measure of L V3 – L SAC connection. Bilateral SAC (BA 7) was strongly associated with well-being according to the previous fMRI (W. Sato et al., 2019) and PET studies (Volkow et al., 2011). SAC is a wide region and one of the sub-regions is precuneus that is strongly related to well-being (Kong, Ding, et al., 2015; W. Sato et al., 2015; W. Sato et al., 2019; Volkow et al., 2011). Precuneus plays an important role in self-awareness (Kjaer et al., 2002), self-consciousness (see review (Cavanna & Trimble, 2006)) and self-reflection (Johnson et al., 2009; Johnson et al., 2006). Previous studies suggested that it combines external and internal information such as personal experience, past memories and future thoughts and this feature might lead precuneus to play an important role in well-being (W. Sato et al., 2015).

## 5. Limitations

In the current study, we used rs-fNIRS to show its efficiency by utilizing two different connectivity measure for classification of flourishing levels and to understand cortical structures that are related to well-being. However, there are some critical limitations that need to be addressed.

First, the number of subjects that participated in this study is low to perform a machine learning study. In machine learning applications, training and testing the model using larger samples allows us to create more robust and reliable models. To overcome this problem, we used nested cross-validation due to producing robust and unbiased estimation without considering the sample size (Vabalas et al., 2019) and also prevented overfitting (Hosseini et al., 2020). However, more extensive research by recruiting larger cohorts ideally from different centers must be realized to increase the reliability of the model and validate the suggested biomarkers.

Second, due to penetration depth limitation of fNIRS which is around 1-2cm depending on source detector distance, it was not possible to measure the sub cortical regions. Used connections was estimated by using only cortical regions. Therefore, we could not observe some direct connections across regions.

Finally, as we stated in methods section, there was no short source-detector separation channels in the dataset to eliminate systemic blood flow on scalp. To remove this effect using short separation channels is a necessity. Due to not being able to perform this, this effect might be available in time series.

## 6. Conclusion

In this study, it has been shown that rs-fNIRS data might be used to discriminate the individuals according to their flourishing levels by utilizing ML techniques. To our best knowledge, this is the first study that utilizes rsfNIRS and ML techniques to classify the individuals according to their flourishing levels. Previous studies utilized some features to understand well-being of individuals by either self-report data or physiological or laboratory data. We found 90% accuracy with 0.90 and 0.93 AUC by using Nearest neighbour and SVM algorithms. This result outperformed the results that were previously reported and showed the efficiency of rs-fNIRS. Also, we also showed that sub-components of DMN in temporo-parieto-occipital region might be potential biomarkers to understand the flourishing levels of individuals. Using fNIRS to reveal these biomarkers is critical, compared to using fMRI due to being inexpensive and having high degree of mobility. Also, rather than using statistical approaches that gives an answer in group level, ML provides us an answer in individual level. However, more extensive research is needed to understand the underlying neural mechanisms of well-being and validate the suggested biomarkers.

## Code availability

The code used for analysis can be provided upon request.

